# Conflicting Auxin-Phosphate Signals Impact on RSL2 Expression and ROS-Homeostasis Linked to Root Hair Growth in *Arabidopsis*

**DOI:** 10.1101/244939

**Authors:** Silvina Mangano, Silvina Paola Denita-Juarez, Eliana Marzol, Cecilia Borassi, José M. Estevez

## Abstract

Here, we examined by which mechanism root hairs integrate conflicting growth-signals like the repressive high Pi-level clue and a concomitant high auxin exposure that should promote growth and questioned if these complex signals might activate known molecular players in polar growth.

---

How plant cells regulate their size is one of the most fascinating questions in current plant biology. Root hairs are single plant cells that can expand several hundred-fold their original size and they are an excellent model system for learning about cell size regulation. Root hair size determines the surface area/volume ratio of the whole roots exposed to nutrient and water pools, thereby likely impacting nutrient and water uptake rates. The speed at which they grow is determined by cell-intrinsic factors like hormones (e.g., auxin and ethylene) and external environmental signals like nutrients availability in the soil (e.g., phosphate and nitrates) (Feng et al. 2017; Marzol et al. 2017). The root hair polar growth program is initially triggered by the transcription factors (TFs) of basic helix-loop-helix (bHLH) family RHD6 (ROOT HAIR DEFECTIVE 6) /RSL1 (ROOT HAIR DEFECTIVE 6 LIKE 1) in the initiation phase, and then, activated by the expression of RSL4/RSL2 during the elongation phase (Menand et al. 2007; Yi et al. 2010). Hormones and environmental cues converge to regulate the expression of RSL4, which controls the final root hair cell size (Yi et al. 2011; Lee et al.2013; Datta el al. 2015; Song et al. 2016; Rymen et al. 2017). Although, in this process, other TFs might act in a coordinative manner under specific signals such as Pi-starvation and ethylene as this is the case for EIN3 (ETHYLENE INSENSITIVE 3)/EIL1 (ETHYLENE INSENSITIVE 3-LIKE 1) and LRL3 (LOTUS JAPONICA ROOTHAIRLESS-LIKE1) (Yi et al. 2010; Song et al. 2016, Feng et al. 2017). Recently, it was found that auxin, by releasing several ARFs (e.g., ARF5, ARF7, ARF8 and ARF19) from Aux/IAA proteins, directly activates RSL4 expression and controls root hair growth linked to ROS-homeostasis involving three RESPIRATORY BURST OXIDASE HOMOLOG proteins (e.g., RBOHC,H,J) and four type-III secreted peroxidases (e.g., PER1,44,73) (Mangano et al. 2017). In addition, RSL2 was also involved in the auxin-mediated growth response although it was unclear its mode of action at the molecular level (Mangano et al. 2017). Conversely, high levels of inorganic phosphates (Pi) in the soil (or in the media) are able to strongly repress RSL4 expression linked to polar growth by an unknown mechanism while Pi-starvation results in an extensive outgrowth of root hairs. This response is associated with increased activity of two MYB-transcription factors, PHR1 (PHOSPHATE STARVATION RESPONSE) and PHR1-LIKE1 (PHL1), the homeodomain transcription factor AL6/PER2 (ALFIN-LIKE6/Pi DEFICIENCY ROOT HAIR DEFECTIVE 2), EIN3/ENL1, and RSL4 (Bustos et al., 2010; Yi et al., 2010; Chandrika et al., 2013; Datta el al. 2015; Song et al. 2016; Feng et al. 2017). Here, we examined by which mechanism root hairs integrate conflicting growth-signals like the repressive high Pi-level clue and a concomitant high auxin exposure that should promote growth and questioned if these complex signals might activate known molecular players in polar growth.

## Auxin overcome Pi-repression of root hair growth in a ROS-dependent manner

Increased levels of Pi in the media (from 1–20 mM) were able to strongly repress root hair growth as previously reported (Datta et al. 2015; Song et al. 2016) as well as to greatly reduce the Reactive Oxygen Species (ROS) levels in root hair cells in the *Arabidopsis thaliana* model. ROS levels were measured at the root hair cell tip with the cell permeable probe H_2_DCF-DA (2’,7’-dichlorodihydrofluorescein diacetate) that becomes irreversibly fluorescent under ROS-oxidation. Then, an exogenous auxin supply (100 nM IAA, Indole 3-Acetic Acid) was able to restore root hair growth independently of the level of Pi in the media (**Figure 1A**). In addition, ROS levels were also re-established in the auxin-treated root hairs even in the presence of high-levels of Pi suggesting that both signals, high-Pi levels as well as high-auxin, operated in an opposed manner by a ROS-dependent mechanism (**Figure 1A**). Then, it was tested whether the suppression of ROS-derived from RBOHs activities with VAS2870 (VAS, a specific RBOHs inhibitor) might affect auxin growth-effect in the presence of Pi (**Figure 1B**). VAS treatment was able to revert the auxin growth-effect at the root hair level by down-regulating RBOHs-derived ROS production (**Figure 1B**). A similar repressive effect was obtained when *rbohc* mutant with low-ROS, was incubated with auxin in the presence of high Pi-levels (**Figure 1B**). This confirms that drastic changes in ROS-homeostasis controlled mostly by RBOHC inhibit auxin growth promoting effect. It was previously shown that the over-expression of either RSL4 or Auxin Response Factor 5 (ARF5) under the control of the root hair EXPANSIN 7 promoter (E7) led to high-levels of ROS (Mangano et al. 2017). In addition, E7:RSL4 and E7:ARF5 were more insensitive to high Pi levels that Wt Col-0 and they developed almost regular extended root hairs (**Figure 1C**). This result indicates that high expression of RSL4 or constitutive activated auxin signaling, both are able to partially repress the high levels of Pi-growth effect. To corroborate the ROS measurements performed with the H_2_DCF-DA probe as well as to determine if the main ROS molecule involved was hydrogen peroxide (H_2_O_2_), we measured cytoplasmic H_2_O_2_ (_cyt_H_2_O_2_) levels using a genetically encoded YFP-based H_2_O_2_ sensor, HyPer (Mangano et al. 2017). Similar results were obtained for high-levels of Pi that first repressed cytoplasmic H_2_O_2_ while exogenous auxin was able to trigger an up-regulation of the ROS-signal in the presence of high-Pi in the root hairs (**Figure 1D**). All together indicates that auxin is able to overcome Pi-growth repression and it requires ROS-production to trigger proper polar-growth.

**Figure 1.**
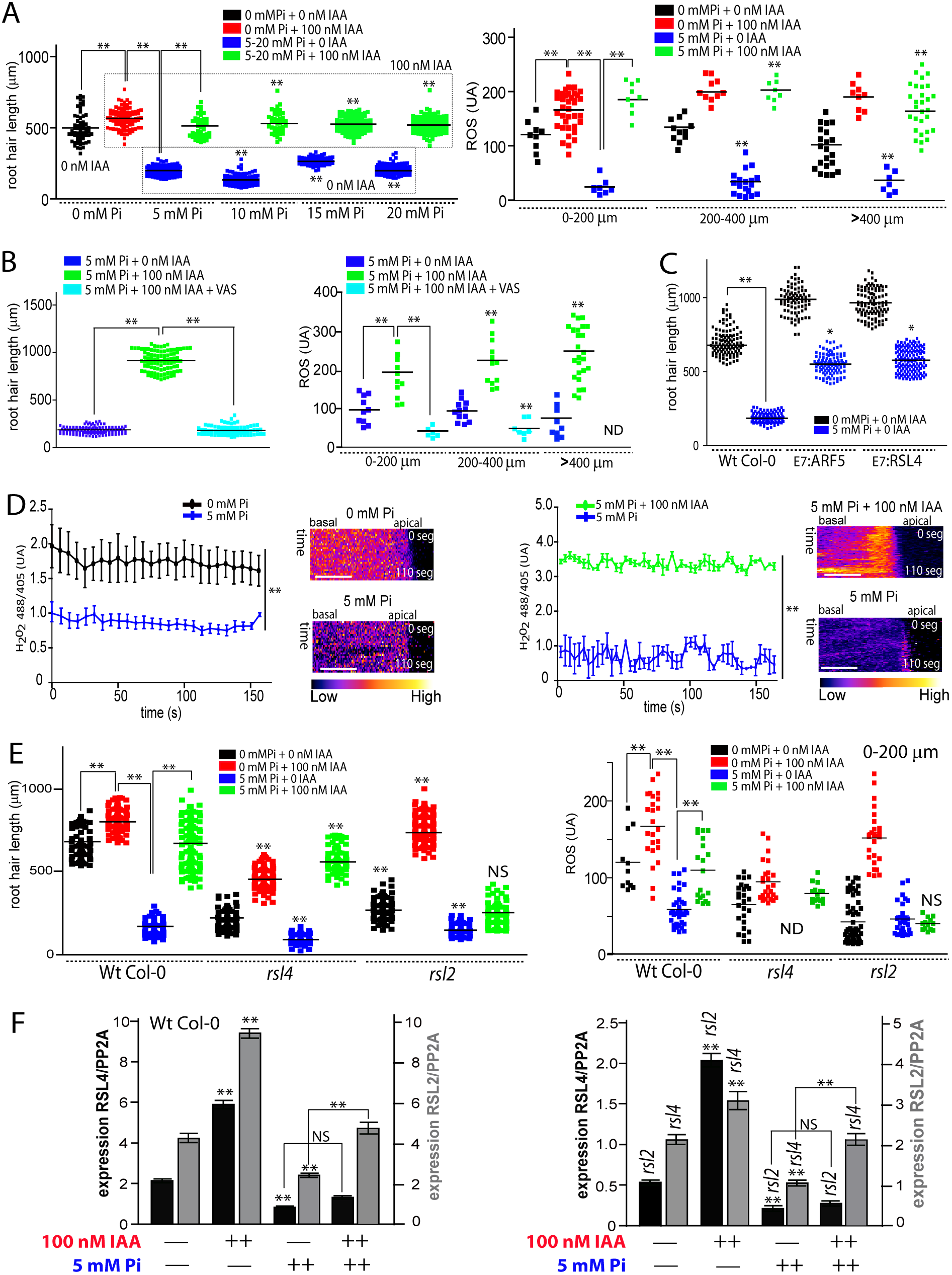
Auxin overcomes high-Pi root hair growth-repression by enhancing RSL2 expression linked to ROS-homeostasis. A. Auxin circumvents root hair growth repression imposed by increased levels of Pi by enhancing ROS-production. Root hair length (mean ± SD) was measured in Wt Col-0 roots non-treated, treated with 100nM IAA (indole 3-acetic acid) or with 5mM inorganic Pi, or both treatments at the same time (on the left). Total ROS levels generated by oxidation of H_2_DCF-DA were measured at the root hair tip in different stages of root hair development in the four different treatments (on the right). B. Auxin recovery of root hair growth is dependent on ROS-production. Root hairs treated with 100 nM IAA in the presence of high levels of Pi and VAS inhibitor fails to recover its normal growth (on the left). ROS-levels were down regulated in the treated root hairs with 100 nM IAA in the presence of high levels of Pi and VAS inhibitor (on the right). ND= root hairs not detected. C. High levels of ARF5 and RSL4 expression in root hairs cells (EXPANSIN 7 promoter, E7) are able to partially overcome the growth repression imposed by high levels of Pi on cell elongation. Comparisons are made between Wt Col-0 and _E7_:ARF5 or _E7_:RSL4 lines under high levels of Pi. D. Auxin trigger changes in _cyt_H_2_O_2_ levels detected with HyPer. _cyt_H_2_O_2_ levels Wt Col-0 root hairs expressing HyPer sensor treated with 5 mM Pi and 100 nM IAA. _cyt_H_2_O_2_ levels are based on the ratio 488/405 nm of HyPer biosensor at the root hair tip over 200 s. On the right, selected kymographs resulting of this analysis only for root hairs of >200 µm in length. Scale bar = 5 µm. E. RSL2 but not RSL4 mediates the auxin recovery response in the presence of high Pi-levels. Root hair length (mean ± SD) was measured in Wt Col-0, *rsl4* and *rsl2* roots non-treated, treated with 100nM IAA (indole 3-acetic acid) or with 5mM inorganic Pi, or both treatments at the same time. ROS-levels were partially recovered in Wt but not in *rsl2* when treated with 100 nM IAA in the presence of high levels of Pi. No ROS was detected in *rsl4* under high levels of Pi. Only root hairs <200 µM in length were analyzed. F. Auxin under high Pi-levels triggers the expression of both, RSL4 and RSL2. Levels of RSL4 and RSL2 expression in Wt Col-0 roots non-treated treated with 100nM IAA (indole 3-acetic acid) or with 5mM inorganic Pi, or both treatments at the same time (on the left). Levels of RSL4 and RSL2 expression in *rsl2* and *rsl4* respectively non-treated treated with 100nM IAA (indole 3-acetic acid) or with 5mM inorganic Pi, or both treatments at the same time (on the left). Each qRT-PCR reaction was performed in triplicate, and each experiment was repeated three times using independent preparations of RNA. Comparisons are made in the same genetic background between levels of gene expression (of RSL2 or RSL4) in non-treated samples (no IAA/no Pi) versus treated with IAA or Pi. In addition, Pi-treated samples are compared only to Pi+IAA-treated roots. In Figs 1A-F, *P*-value of one-way ANOVA, (**) P<0.001, (*) P<0.01. NS= not significant differences. Error bars indicate ±SD from 3 biological replicates.

## Expression of RSL2 is required to bypass Pi-growth repression in the presence of auxin

In order to have insights on the molecular mechanism behind these high Pi-auxin conflicting responses, we tested root hair growth linked to ROS-levels in the absence of RSL2 and RSL4 TFs that were shown to regulate the auxin-mediated root hair growth-response (Datta et al. 2015; Mangano et al. 2017). The double mutant *rsl2 rsl4* is not capable of develop visible root hairs in any condition tested, either under high levels of auxin or ethylene or nutrient deprivation suggesting that both TFs are necessary to run the basic transcriptional machinery to produce root hairs (Mangano et. al 2017; Feng et al. 2017). High Pi-level strongly represses the expression of both, RSL2 and RSL4 TFs in the Wt Col-0 roots while auxin treatment up-regulates both TFs, and not only RSL4 as previously indicated (Pires et al. 2013). When *rsl4* (that lacks RSL4 transcripts) was tested, the response was similar to Wt Col-0 root hairs but strongly attenuated indicating that auxin can rescue the negative effect of high-Pi levels, possibly mediated by RSL2 action. In agreement with this idea, when RSL2 was lacking (in *rsl2* mutant), there was almost no response associated to the auxin treatment (**Figure 1E**). This indicates that RSL2 mediates the auxin rescue-response in the presence of high Pi-levels. In addition, ROS measurements in Wt Col-0 as well as in *rsl4* and *rsl2* mutants positively correlate with growth responses in all conditions tested (**Figure 1F**). As a complementary approach, levels of RSL2 and RSL4 transcripts were measured in their respective mutant backgrounds *(rsl4* and *rsl2*, respectively) to define if any of these two TFs depends on the presence of the other to be expressed under these conflicting growth conditions (**Figure 1F**). In the *rsl4* mutant background, the expression of RSL2 was down regulated 50% to the levels found in Wt Col-0, while, in *rsl2* background, the level of RSL4 expression was three to four-times lower in all conditions measured (e.g., no Pi or auxin added, high Pi-level, high auxin, or both). This indicates that the absence of RSL2 might affect RSL4 expression, and when RSL4 is lacking, RSL2 transcripts also decrease suggesting a positive feed forward loop between RSL4-RSL2 by a direct or indirect mechanism that requires further investigation.

Several possible scenarios can now be considered to understand how auxin together with high Pi-levels control root hair growth (**Figure S2**). Since high levels of Pi strongly repress root hair growth and ROS-production, Pi might operate at several regulatory points such as down regulating auxin biosynthesis, auxin-conjugation and transport within the roots towards the trichoblast cells (Velasquez et al. 2016), and then, indirectly affecting RSL4/RSL2 expression. Alternatively, high Pi levels could directly repress RSL4/RSL2 expression by an unknown mechanism. Recently, abscisic acid (ABA) throughout the TF OBP4 (OBF BINDING PROTEIN 4) was able to repress RSL2 (and RSL3) expression indicating that multiple hormonal signals on top of auxin may operate in these cells as previously described (Rymen et al. 2017). In addition, it is plausible that RSL2 and RSL4 would be able to form protein dimers as reported in other several TFs that contains HLH domain (Feller et al. 2011), more precisely heterodimers (e.g., RSL2-RSL4) and homodimers (e.g., RSL2-RSL2 and RSL4-RSL4) with slightly different transcriptional downstream target genes. Since it has been shown that RSL4 is able to self-activate by a forward positive-loop (Hwang et al. 2017), we hypothesized that RSL2 would be also capable to the same. In addition, RSL2 would be able to up-regulate RSL4, and vice versa as it was suggested in a previous work (Pires et al. 2013). Further studies are now required to establish the interconnections between both TFs under the different conditions of auxin and Pi tested in this work. Second, since RSL4 is directly activated by ARF5 (and possible several other ARFs including ARF7,8,19) in trichoblast cells (Mangano et. al 2017), we also postulate that RSL2 would be up-regulated by some of these ARFs since auxin triggers its expression two and half-times (**Figure 1E**). In agreement, three consensus Aux-RE sites are found in the regulatory region of RSL2 as possible targets for ARFs binding (not shown). Still how the fine-tune regulation of ARFs on the RSL2-RSL4 is coordinated remains to be discovered. In summary, although it has been shown before that auxin triggers root hair cell elongation while high Pi-levels repress its growth, it was unknown how both conflicting signals might act together. This study identifies a new layer of complexity between RSL2/RSL4 TFs acting on ROS-homeostasis under conflicting growth-signals at the root hair level.

## Acknowledgements

We apologize for the inadvertent omission of any pertinent original references owing to space limitations. No conflict of interest is declared.

## Author’s contributions

S.M performed all the experiments, reviewed the text, figures, and references; S.P.D.J. performed some of the HyPer experiments; E.M. and C.B. reviewed text, references, and figures; J.M.E conceived the project, designed the figures, and wrote the article with contributions of all the authors.

## Financial source

This work was supported by a grant from ANPCyT (PICT2014–0504 and PICT2016–0132) and ICGEB (CRP/ARG16–03 Grant) to J.M.E and a grant from ANPCyT (PICT2016–0491) to S.M.

## Supplementary Information

Material and Methods. Supplementary Figures S1-S2. Supplementary Tables S1-S2.

### Materials and Methods.

**Plant Growth and mutant isolation**. *Arabidopsis thaliana* Columbia-0 (Col-0) was used as the wild type (Wt) genotype in all experiments, unless stated otherwise. All mutants and transgenic lines tested are in this ecotype. Seedlings were germinated on agar plates in a Percival incubator at 22°C in a growth room with 16h light/8h dark cycles for 10 days at 140 µmol m^−2^s^−1^ light intensity. Plants were transferred to soil for growth under the same conditions as previously described at 22°C. For identification of T-DNA knockout lines, genomic DNA was extracted from rosette leaves. Confirmation by PCR of a single and multiple T-DNA insertions in the target RBOH and PER genes were performed using an insertion-specific LBb1 or LBb1.3 (for SALK lines) or Lb3 (for SAIL lines) primer in addition to one gene-specific primer. To ensure gene disruptions, PCR was also run using two gene-specific primers, expecting bands corresponding to fragments larger than in Wt. In this way, we isolated homozygous lines (for all the genes mentioned above). Mutant list is detailed in **Supplementary Table S1**.

**Growth media. RBOHs inhibition and auxin treatment**. Sterilized seeds were stored at 4°C in sterile water were for 48 hours and then were germinated on agar plates containing modified Hoagland solution contained 1 mM Ca(NO_3_)_2_, 50 µM CaCl_2_, 0.25 mM MgSO_4_, 50 µM Fe-NaEDTA, 1 mM KCl, 2 µM MnSO_2_–2H_2_O, 0.5 µM CuSO_4_–5H_2_O, 0.5 µM Na_2_MoO_4_–2H_2_O, 2 µM Zn SO_4_–7H_2_O and 2.5mM MES containing low phosphate (5 µM (NH_4_)2HPO_4_). Media were adjusted to a pH of 5.7 and solidified using 0.8% agar. After 5 days plants were transferred into to modified Hoagland solution with low phosphate (5 µM (NH_4_)2HPO_4_) or high phosphate (5 mM (NH_4_)_2_HPO_4_). Low and high phosphate were combined with 100 nM of indole-3-acetic acid (IAA) or/and 15 µM of VAS2870 (VAS, for 3-Benzyl-7-(2-benzoxazolyl)thio-1,2,3-triazolo(4,5-d)pyrimidine). After 5 days (10 days old), quantitative analysis of root hair phenotypes and total ROS measurements with H_2_DCF-DA were made.

**Root hair phenotype**. For quantitative analysis of root hair phenotypes in *rboh* mutants and Wt Col-0, 200 fully elongated root hairs were measured (n roots= 30) from seedlings grown on vertical plates for 10 days. Values are reported as the mean ±SD using the Image J 1.50b software. Measurements were made on images were captured with an Olympus SZX7 Zoom microscope equipped with a Q-Colors digital camera.

**H_2_DCF-DA probe used to measure total ROS**. Arabidopsis seeds were grown in plates with sterile agar 1% for 8 days in chamber at 22°C with continuous light. These seedlings were incubated with 2’,7’-Dichlorodihydrofluorescein diacetate (H_2_DCF-DA) in darkness in a slide for 10 min 50 µM at room temperature. Samples were observed with a confocal microscope equipped with 488 nm argon laser and BA510IF filter sets. It was used the following configuration (10X objective, 0.30 NA; 4.7 laser intensity, 1.1 off set, 440 PMT for highest ROS levels and 480 PMT for ROS media and 3 of gain. Images were taken scanning XZY with a 2 um between focal planes. Images were analyzed using ImageJ. To measure ROS highest levels, a circular ROI (r=2.5) was taken in the zone of the root hair with highest intensities. To measure ROS mean the total area of the root hair was taken. Pharmacological treatments were carried out with a combination of the following reagents: 1–20 mM Pi, 100 nM IAA, and 15 µM of VAS2870. The sample was washed with a MS 0.5X solution and the image acquisition was made with 10X objective and 400ms of exposure-time in an epifluorescence microscope (Zeiss, Imager A2). To measure ROS levels a circular ROI (r=2.5) was taken in the tip of the root hair. Values are reported as the mean ±SD using the Image J 1.50b software.

**HyPer sensor to measure _cyt_H_2_O_2_**. HyPer consists of a circularly permuted YFP (cpYFP) molecule coupled to a regulatory domain of the *Escherichia coli* H_2_O_2_ sensor OxyR (1–4). When exposed to H_2_O_2_, the excitation peak of cpYFP shifts from 420 to 500 nm, while the emission peak remains at 516 nm allowing it to be used as a ratiometric biosensor (Mangano et al. 2017). Ten day-old *Arabidopsis* seedlings expressing the fluorescent HyPer biosensor were used. Root hairs were ratio imaged with the Zeiss LSM 510 laser scanning confocal microscope (Carl Zeiss) using a 40X oil-immersion, 1.2 numerical aperture. The HyPer biosensor was excited with both the 405 nm blue diode laser and with the 488 nm argon laser. The emission (516 nm) was collected using a primary dichroic mirror and the Meta-detector of the microscope. For time-lapse analysis, images were collected every 3s. To measure ROS highest levels, a circular ROI (r=2.5) was taken in the root hair tip for each image the time lapse. Treatments were made *in vivo* with 5mM Pi, 100 nM IAA, or 50 µM of VAS2870. Values are reported as the mean ±SD using the Image J 1.50b software.

**Quantitative reverse transcriptase PCR (qRT-PCR)**. Total RNA was isolated from 10-d-old seedling roots (40 for each line) using the RNAzol RT (MRC). cDNA was synthesized using M-MLV Reverse Transcriptase (Promega). qRT-PCR analyses were performed using LightCycler480 SYBR Green I Master-Roche. Gene-specific signals were normalized relatively to PP2A (AT1G69960; serine/threonine protein phosphatase 2A) signals. Each qRT-PCR reaction was performed in triplicate, and each experiment was repeated three times using independent preparations of RNA. Primers used are as listed in **Supplementary Table S2**.

### Legend to Figure S1.

**Figure S1.**
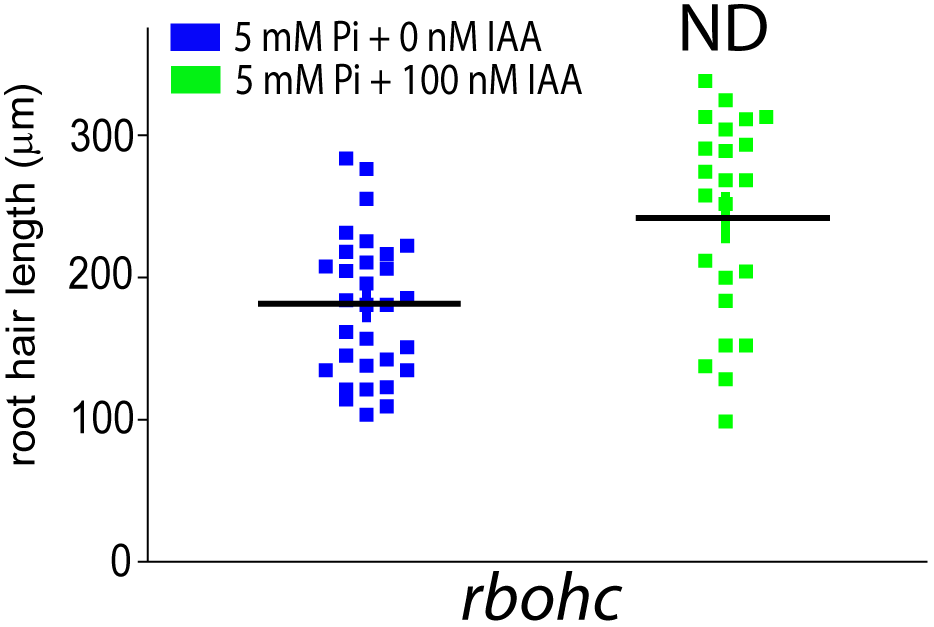
Auxin fails to overcome high-Pi root hair growth-repression in a low ROS-depleted background (in *noxc* and *per44,73* mutants). Auxin fails to circumvent root hair growth repression imposed by increased levels of Pi in a ROS-depleted *rbohc* mutant. Root hair length (mean ± SD) was measured in *rbohc* roots treated with with 5mM inorganic Pi with or without 100nM IAA (indole 3-acetic acid) (on the left). Total ROS levels generated by oxidation of H_2_DCF-DA were measured at the root hair tip in different stages of root hair development in the four different treatments (on the right).

**Figure S2.**
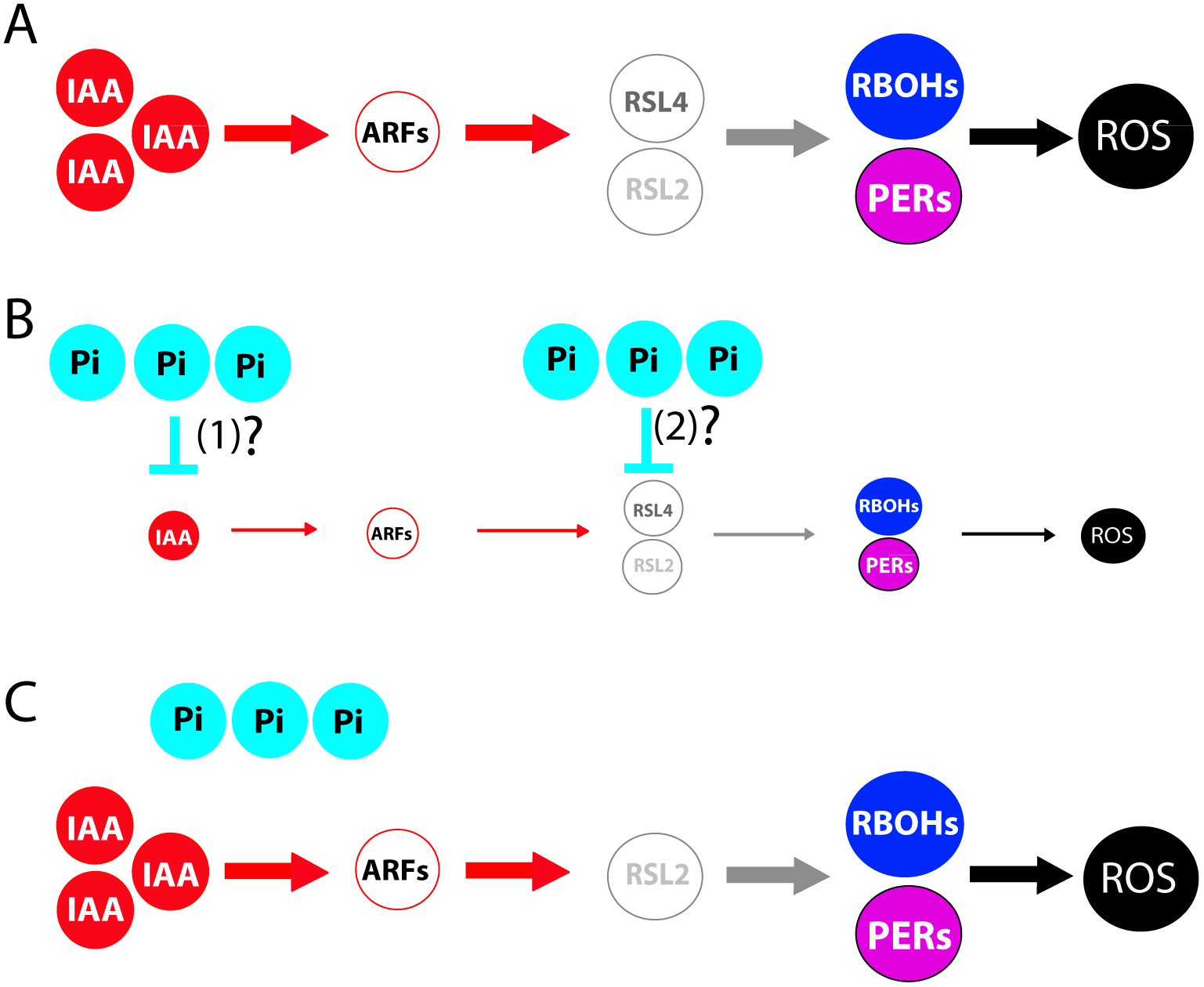
Proposed models of auxin-RSL2/RSL4 regulation of ROS mediated polar root hair growth in the absence and in presence of high-Pi. Transcriptional responses under high-level of auxin in the absence of Pi (A), in the presence of high Pi with very low-level of auxin (B), and in high levels of Pi together with high levels of auxin (C) are shown. A. The bHLH transcription factor RSL4 as well as RSL2 are both transcriptionally activated by high levels of auxin (IAA) and its expression is directly regulated by several Auxin Response Factors (ARFs; ARF5,7,8,19). Throughout RSL4 (and possibly RSL2), auxin activates the expression of two RBOHs (RBOHC,J) and four PERs (PER1,44,60,73) that together regulate ROS homeostasis in the apoplast (in combination with RBOHH). Based on Mangano et al. (2017) and the present work. B. High level of Pi in a low auxin level environment would be sufficient to repress the RSL4/RSL2 transcriptional response. Pi repression act depleting auxin levels in the atricoblast (1) or repressing by an unknown mechanism directly the expression of RSL4/RSL2 (2). C. High levels of Pi in the presence of high levels of auxin are able to activate the expression of RSL2 and control ROS-homeostasis derived from RBOHs and PERs activities. Solid lines indicate transcriptional activation or metabolite production. Reactive Oxygen Species (ROS) include hydroxyl radical (^•^OH), superoxide ion (O_2_^•−^) and hydrogen peroxide (H_2_O_2_).

**Supplementary Table S1.**
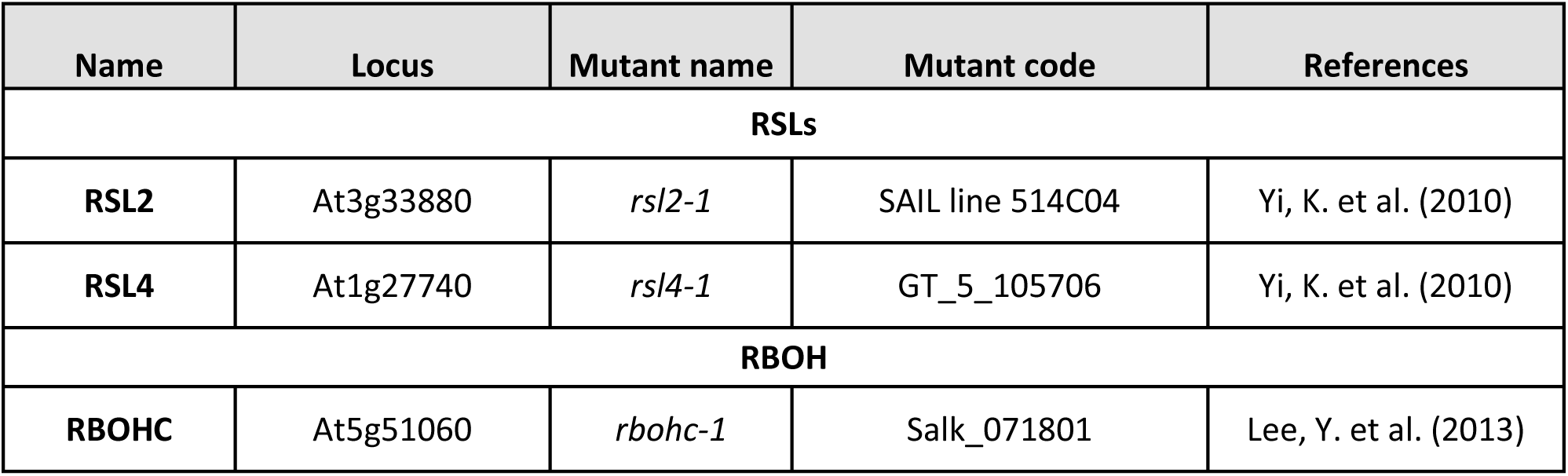
RSLs and NADPH oxidase C (RBOHC) mutant line used in this study. All are in Col-0 background.

**Supplementary Table S2.**
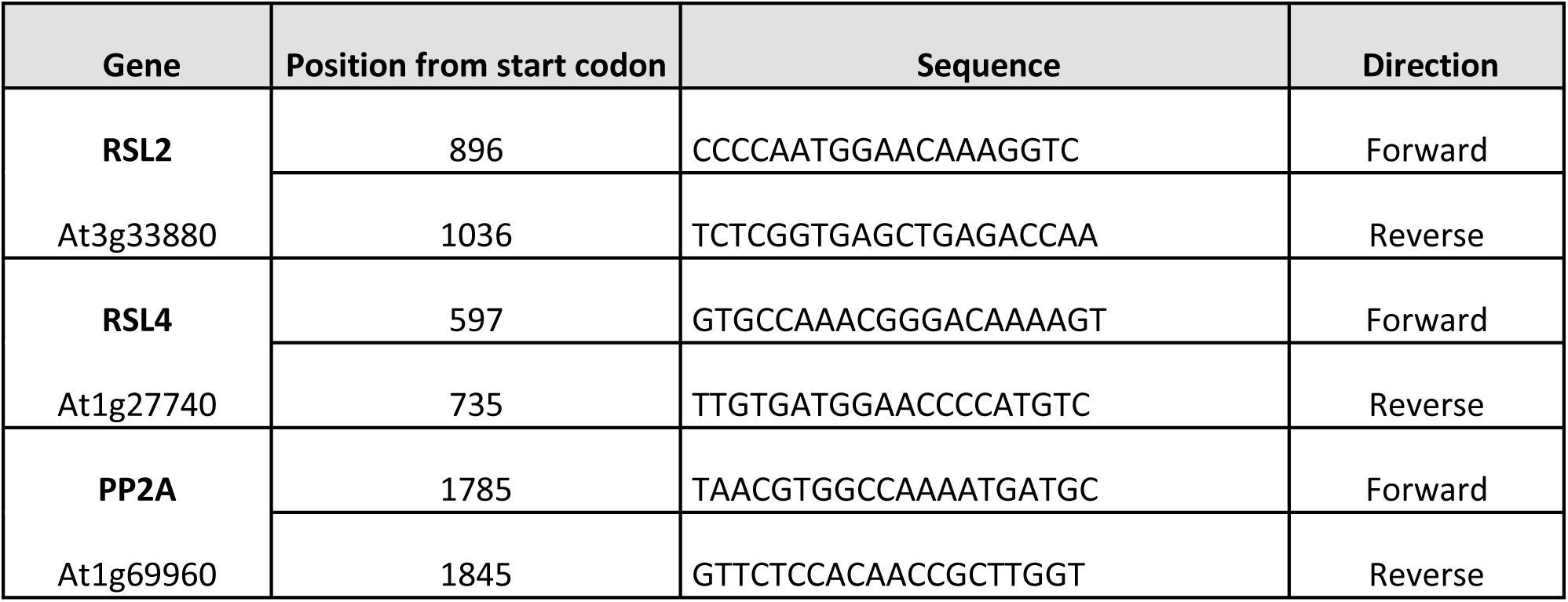
Primer List used for qPCR.

